# Chasing the fitness optimum: temporal variation in the genetic and environmental expression of life-history traits for a perennial plant

**DOI:** 10.1101/2021.10.12.464067

**Authors:** Mason W. Kulbaba, Zebadiah Yoko, Jill A. Hamilton

**Author notes:** Author for Correspondence: Jill Hamilton, Department of Ecosystem Science and Management, 323 Forest Resources Building, University Park, PA.

## Abstract

The ability of plants to track shifting fitness optima is crucial within the context of global change, where increasing environmental extremes may have dramatic consequences to life history, fitness, and ultimately species persistence. However, to track changing conditions relies upon the complex relationship between genetic and environmental variance, where selection may favor plasticity, the evolution of genetic differences, or both depending on the spatial and temporal scale of environmental heterogeneity. Over three years, we compared the genetic and environmental components of phenological and life-history variation in a common environment for the spring perennial *Geum triflorum*. Populations were sourced from alvar habitats that exhibit extreme, but predictable annual flood-desiccation cycles and prairie habitats that exhibit similar, but less predictable variation in water availability. Narrow-sense heritabilities were generally higher for early life history (emergence probability) relative to later life history traits (total seed mass), indicating that traits associated with establishment within an environment are under stronger genetic control relative to later life-history fitness expressions, where plasticity may play a larger role. This pattern was particularly notable in seeds sourced from environmentally extreme, but predictable alvar habitats relative to less predictable prairie seed sources. Fitness landscapes based on seed source origin, largely characterized by varying water availability and flower production, described selection as the degree of maladaptation to the prairie common garden environment relative to seed source environment. Plants from alvar populations were consistently closer to the fitness optimum across all years. Annually, the breadth of the fitness optimum expanded primarily along a moisture gradient, with inclusion of more populations onto the expanding optimum. These results highlight the importance of temporally and spatially varying selection for the evolution of life history, indicating plasticity within perennial systems may over time become the primary mechanism to track fitness for later life history events.

## Introduction

Environmental cues essential to life history transitions may shift as a consequence of global change, increasing the potential discrepancy between realized and optimal fitness (Reed *et al*, 2010; Edelaar *et al*, 2017). Fitness convergence requires a combination of strategies, including the evolution of locally adapted genotypes that are the product of natural selection (Clausen *et al*, 1941; Hereford, 2009; Kawecki and Ebert, 2004; Leimu and Fischer, 2008) and phenotypic plasticity, the ability of individual genotypes to directly alter phenotypes in response to the environment (Bradshaw, 1965; Scheiner, 1993; Josephs, 2018). As environments change, ensuring individuals have the capacity to maximize fitness through these strategies is crucial to population persistence. However, the relative degree to which adaptive evolution or plasticity evolves can vary across traits, particularly if different life history stages disproportionately mediate the fitness effects of environmental heterogeneity (Roff and Mousseau, 1987). Climate models predict increasing environmental heterogeneity and decreased temporal and spatial predictability of environmental cues that may directly influence the ability of individuals to track fitness optima via plasticity or genetic adaptation (Anderson and Gezon, 2015; Baythavong, 2011; Franks *et al*, 2013). Consequently, within the context of climate change it is essential to understand the impact that spatial and temporally varying environments may have to adaptive genetic differentiation and plasticity for life-history traits and long-term fitness expressions.

Realized phenotypes are the combined products of genetic and environmental effects and fitness is an emergent property determined by the degree of similarity between individual phenotypes and environmentally determined optima. With this two-fold contribution, limited variation in heritable effects or plastic responses to environmental conditions can constrain the match between optimal and realized phenotypes, ultimately impacting fitness. The relative amount of phenotypic variation attributed to heritable and plastic components will differ both among traits (Mousseau and Roff, 1987; Price and Schluter, 1991) and environments (Hoffmann and Merila, 1999). For example, phenological schedules require at least some level of genetic determination to initiate life-history events, but plasticity is often required to achieve maximum fitness given variability inherent to natural environments. Theory suggests the proportion of phenotypic variation attributed to plasticity should be greater in environments with repeated and predictable cues compared to more unpredictable environments (Lande, 2009; Botero *et al*, 2015; Tufto, 2015; Leung *et al*, 2020). However, plasticity may be constrained when there is limited capacity (Auld *et al*, 2010) or the maintenance and expression of plasticity is costly (DeWitt *et al*, 1998). Greater plasticity may also evolve in traits closely linked to fitness to both maintain fitness under unfavorable conditions and maximize fitness when conditions are more favorable (Sultan, 2001). With expected increases in environmental variability at a global scale and decreased predictability of cues under climate change (Botero *et al*, 2015), the likelihood of potential mismatches between phenotypes and environments is increasing. This trend highlights the need to refine our understanding of variation in the genetic and environmental components contributing to life history evolution across spatially and temporally varying environments.

A major challenge to predicting responses to environmental change is the complexity of interacting effects. This is particularly evident where exposure to varying degrees of micro-environmental variation influence a populations’ adaptive capacity (Denney *et al*, 2020). Quantitative genetic theory predicts that trait heritability varies across environments and time (Lynch and Walsh, 1998; Price and Schluter, 1991) and environmental heterogeneity can contribute to the evolution of plasticity across a species’ range (Baythavong, 2011; Edelaar *et al*, 2017; Kingsolver *et al*, 2002; Scheiner, 2013). Selection for plasticity should occur in environments with predictable cues, such as those marking seasonal transitions. Such consistent selection may result in reduced genetic variation for plasticity (Oostra *et al*, 2018). This contrasts with expectations for populations sourced from unpredictable environments, where genetic variation in plastic responses is expected to be maintained, enhancing trait plasticity (Ghalambor *et al*, 2007; Chevin *et al*, 2010;). Given these contrasting predictions, understanding the degree to which environmental heterogeneity and plasticity interact to influence phenotypic variation is essential to predicting population fitness over space and time. However, quantifying adaptive evolution requires detailed estimates of the degree of constancy for quantitative genetic parameters across time and environments (Young *et al*, 1994; Bemmels and Anderson, 2019). Quantifying the relative influence of genetic and environmental variation to life history traits in natural settings is difficult as the consistency of environmental cues cannot be experimentally controlled. Thus, common garden experiments leveraging ecotypic variation present experimental designs necessary to facilitate estimation of quantitative parameters across space and time.

In the present study, we test the prediction that individuals sourced from predictably heterogeneous environments will exhibit greater heritability for life history traits relative to those from less predictable, but still heterogeneous environments. In addition to environment-specific variation in heritability, we expect that the heritability of traits varies with ontogeny. Specifically, we predict that early life history traits, such as emergence, will be under greater genetic control relative to later life history stages, representing a continuum of relatively heritable to plastic traits. Few studies have quantified the relative genetic and environmental contributions to life history in perennial plant species due to challenges associated with estimating lifetime fitness (but see Campbell, 1996; Campbell, 1997; Simons and Johnston, 2000). However, these data remain key to quantifying the capacity of long-lived species to traverse temporal and spatially varying fitness landscapes. Finally, we evaluate individuals’ ability to traverse the fitness landscape following an experimental increase in distance between a home and novel common garden environment. Quantifying the impact environmental predictability may have to the evolution of life history, and the ability of species to compensate for mismatches across life history stages will be essential to predicting the capacity for species to track changing conditions and modify life-history strategies under global change.

We quantified temporal and spatial genetic variances for traits associated with life history and fitness within the perennial forb *Geum triflorum* Pursch. sourced from distinct habitats with contrasting predictability of environmental cues. *G. triflorum* is an early-season, spring perennial common to midwestern prairie habitats generally characterized by cold, dry winters and hot, humid summers that experience shifts in water availability both annually and seasonally that are relatively unpredictable (Hamilton and Eckert, 2007; Yoko *et al*, 2020). This contrasts with populations of *G. triflorum* persistent on alvar habitats isolated throughout the Great Lakes region of North America. Alvar habitats experience extreme, but predictable annual seasonal variation in water availability from complete flooding in early spring to complete desiccation by early summer (Catling and Brownell, 1995; Stark *et al*, 2004; Yoko *et al*, 2020). Specifically, we ask (i) across a sequential continuum of life history events, is there variance in the heritability of life history traits and expressions of fitness, and if so, are there consequences to lifetime fitness? (ii) Do estimates of evolvability for life-history traits and expressions of fitness indicate the potential for genotypes to become locally adapted to the common garden environment? (iii) Does the heritability of phenology and life history events impede the ability of individuals to maximize proximity to the optimum of the fitness landscape? Understanding how spatially and temporally varying selection interact with heritable genetic and environmental components across species’ life history will be essential to predicting fitness under global change.

## Methods

### Study site and species

This study focuses on *Geum triflorum*, commonly known as Prairie Smoke, a widespread perennial forb from the Rosaceae family. Prairie Smoke is largely distributed across native prairie habitat throughout the Great Plains of North America but is also common on geographically disjunct alvar habitats dispersed around the Great Lakes and into Manitoba, Canada. Open-pollinated maternal seed families were collected in the spring of 2015 from nineteen populations of *G. triflorum*; including eleven populations from the Great Lake alvar region (GLA), two from the Manitoba alvar region (MBA), and six from the prairie (PRA) region of the Midwest (Yoko *et al*, 2020). Within each population approximately 40 seed families were collected along a 100 m transect (as in Hamilton and Eckert, 2007). To supplement these collections, three additional bulk seed collections for prairie populations were obtained from commercial seed providers (SD-PMG, MN-PMG) and the USDA-Pullman Plant Materials Center (WA-BLK) (Fig. 1).

**Figure 1.**
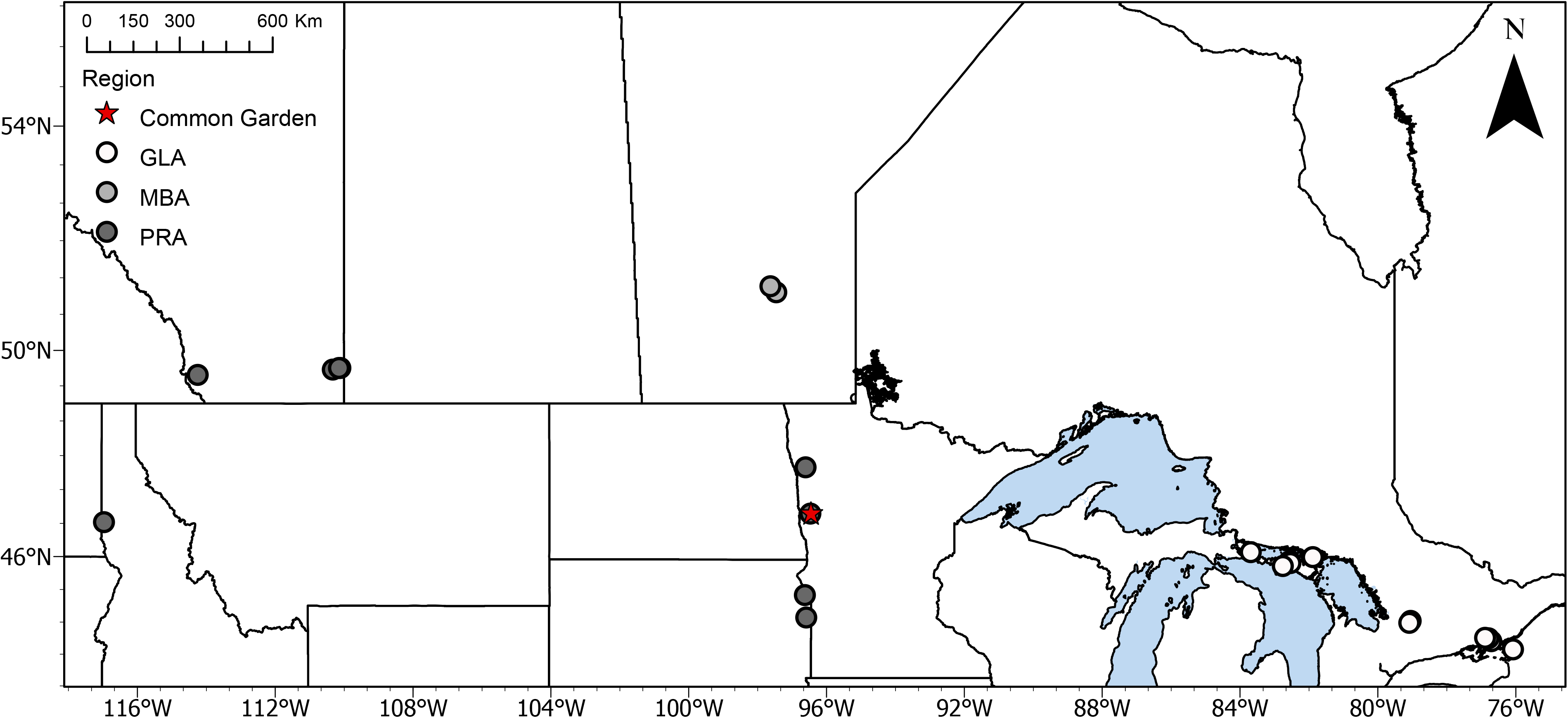
Map indicating location of common garden (star) and Great Lakes Alvar (GLA – open circles), Manitoba Alvars (MBA – light grey circles), and Prairie (PRA – dark grey circles) source populations of *Geum triflorum*.

On November 7, 2015 a common garden experiment was established using open-pollinated seeds at North Dakota State University (described in Yoko *et al* 2020). Using a randomized complete block design, ten maternal seed families were planted for each population across twelve blocks, including 12 individual half-sibs per maternal family. Two replicates were planted per block for bulk-collected seed, for a total of 24 seeds per bulk collection. In total, 2348 individual seeds treated with 0.02% PPM fungicide were planted in ‘cone-tainers’ (158 mL, Stuewe & Sons) filled with Sungro horticulture mix (1N:45P:12K) soil in a greenhouse at North Dakota State University (Table 1). The greenhouse was maintained at 15h days with supplemental daylight from halide lighting at a measured flux density of 0.3383 mmol m2 s-1 for the duration of the experiment and temperatures fluctuating between 18.3°C and 23.9°C. Seedlings were watered bi-weekly and provided a slow release fertilizer mix (Osmocote 14N-14P-14K) throughout the course of the experiment. In May 2016, surviving seedlings were transferred to a permanent outdoor research facility at the Minnesota State University at Moorhead Regional Science Center (46.86913°N, −96.4522°W). Individuals were planted directly into the ground through cut outs in a weed barrier to limit competition. The randomized block design established in the greenhouse was maintained at the permanent field site.

**Table 1.** Source populations of *G. triflorum* collected in 2015 spanning three distinct eco-regions (Great Lake alvars (GLA), Manitoba alvars (MBA), and Prairies (PRA)), along with latitude, longitude, and elevation (m) of population collection sites. Distance from common garden (km) notes the greater circle distance calculated between the population origin and the common garden experiment established at Minnesota State University at Moorhead Regional Science Center, Moorhead, Minnesota, United States.

| Population ID | Region | Latitude | Longitude | Elevation (m) | Distance from Common Garden Experiment (km) |
| --- | --- | --- | --- | --- | --- |
| CAR-NBA | GLA | 44.69 | -79.05 | 268 | 1368 |
| CAR-PSR | GLA | 44.65 | -79.09 | 250 | 1366 |
| MAN-FOX | GLA | 45.90 | -82.58 | 186 | 1068 |
| MAN-KIP | GLA | 45.87 | -82.54 | 183 | 1072 |
| MAN-LCI | GLA | 45.99 | -81.89 | 182 | 1118 |
| MAN-MIS | GLA | 45.81 | -82.76 | 193 | 1056 |
| MI-DRI | GLA | 46.09 | -83.69 | 188 | 980 |
| NAP-ASS | GLA | 44.27 | -76.71 | 126 | 1559 |
| NAP-CE | GLA | 44.33 | -76.79 | 166 | 1551 |
| NAP-SCH | GLA | 44.34 | -76.89 | 154 | 1543 |
| WNY-CB | GLA | 44.10 | -76.08 | 93 | 1613 |
| MB-CRN | MBA | 51.07 | -97.46 | 231 | 473 |
| MB-MR | MBA | 51.18 | -97.63 | 231 | 487 |
| AB-HSC | PRA | 49.64 | -110.33 | 721 | 1071 |
| AB-LL | PRA | 49.54 | 114.25 | 929 | 1348 |
| AB-RL | PRA | 49.67 | -110.11 | 721 | 1056 |
| AB-RO | PRA | 49.67 | 110.15 | 721 | 1059 |
| MN-PMG | PRA | 47.77 | -96.61 | 267 | 101 |
| ND-BSP | PRA | 46.86 | -96.47 | 274 | 2 |
| SD-MUD | PRA | 44.76 | -96.59 | 531 | 234 |
| SD-PMG | PRA | 45.22 | -96.63 | 351 | 184 |
| WA-BLK | PRA | 46.69 | -116.97 | 786 | 1558 |
| CommonGarden Experiment | PRA | 46.87 | -96.45 | 259 | - |

### Data Collection

Phenological and life-history fitness components were evaluated within the common garden from 2015 to the end of the growing season in 2018. Single season phenological observations included the number of days from planting to emergence and establishment of true leaves, recorded in 2015. In addition, multi-year observations were taken for number of days between planting and bolting (defined as the initial elongation of the flowering stem to ~7cm, recorded in 2017 and 2018), days between planting and flowering (recorded annually between 2015-2018) and days between planting and the initiation of infructescence development (defined as the date developing woolly styles extend beyond the corolla to form a diaspore for dispersal, recorded in 2017 and 2018).

To capture annual estimates of reproductive output in the common garden, cloth mesh bags (Uline S-13940) were tied and labeled around each individual infructescence as the woolly styles began to extend beyond the corolla. Cloth mesh bags provide the opportunity for the diaspore to fully mature, while limiting potential loss of reproductive output via wind dispersal. Cloth mesh bags were harvested in each August of the monitoring year. Number of reproductive stems, identified as stems with infructecenses, was used to quantify the number of fruits produced per individual. Seed mass was taken to reflect potential reproductive output per reproductive stem. Total annual fitness was estimated as the cumulative seed mass produced per individual genotype based on all reproductive stems produced. Seed mass is considered a proxy for the number of seeds produced per individual. In 2016, we performed a regression between the number of seeds produced and seed mass for one reproductive stem per individual planted within the common garden (R^2^ = 0.528). Flowering only stems were also quantified in the field as individuals that had flowered, but senesced prior to producing seed, and thus considered non-reproductive. To quantify the total number of flowers produced per individual within each season we combined the total number of reproductive stems with flowering only stems. Those individuals that did not produce a flowering-only stem or reproductive (flower + fruit) stems were noted each year as having survived but were classed as completely non-reproductive.

### Statistical Analyses

To determine the narrow-sense heritability 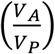 of life-history and phenological traits, we used generalized linear mixed models (Stroup, 2013) as implemented in the package lme4 (Bates *et al*, 2014) using a maternal half-sibling sampling design (Lynch and Walsh, 1998). These analyses employed the most appropriate sampling distributions and link functions for each trait (see Table 2), as determined through comparison of AIC values across models. All analyses included the effect of block and population as fixed effects, maternal family as a random effect, and the interaction between block and maternal family. Following Ahrens *et al* (2020) we calculated family-level heritability as:

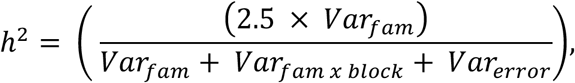

where *Var_fam_* is the maternal family variance, *Var_fam x block_* as the variance attributed to the interaction between family and block, and *Var_error_* as the error/residual variance. Narrow-sense heritability calculations from half-sibling analyses typically multiply the numerator, *Var_fam_*, by 4 to account for half-sibling relatedness (Lynch and Walsh, 1998). However, *G. triflorum* exhibits a mixed mating system that may lead to inbreeding, and the presence of inbred individuals may inflate heritability estimates. Therefore, we follow Ahrens *et al* (2020) and use a factor 2.5 that corresponds to a coefficient of relatedness of r = 1/ 2.5, or approximately a 30% selfing rate. This predicted rate of selfing rate may be high but given that half-sibling analyses represents an upper limit to narrow-sense heritability, our calculation produces a conservative final estimate. Finally, we estimated evolvabilities following Houle (1992) as:

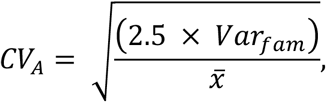

where 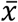 is the trait mean that evolvability is being calculated.

**Table 2.**
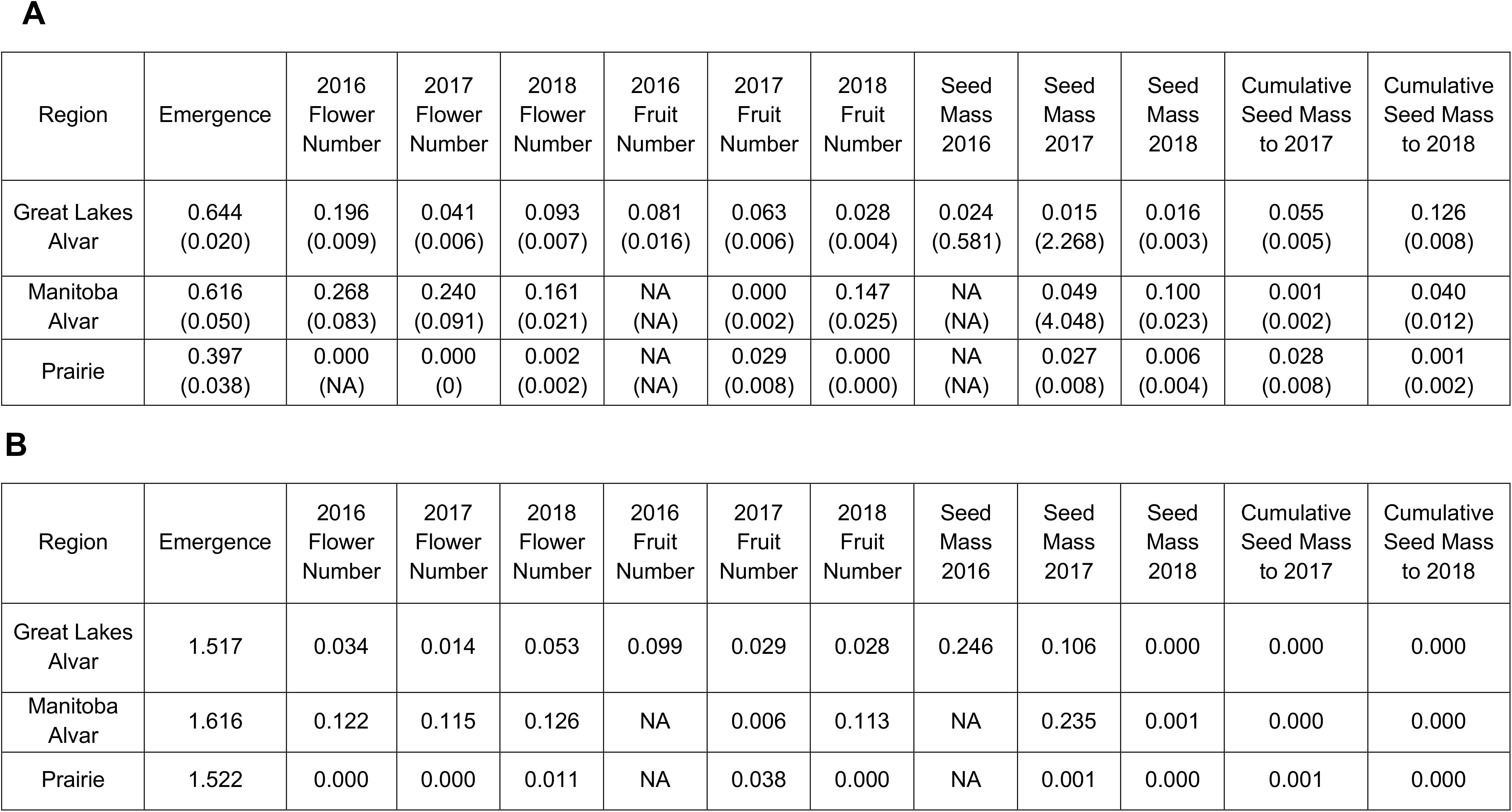
(A) Narrow-sense heritability and standard error and (B) evolvability (coefficient of additive genetic variation) of *Geum triflorum* fitness expressions grown in a common garden environment.

Over the course of our experiment, total fitness was estimated using aster models (Geyer *et al*, 2007; Shaw *et al*, 2008). Mean fitness was estimated for each year of the study for each of the Great Lakes alvars, Manitoba alvars, and prairie regions. The graphical model for fitness (Fig. 2) included individual plant assessments of emergence, survival across years, and survival to flowering modeled as Bernoulli distributions. The production of flowers, fruits, and seed mass were modeled as negative binomial distributions. The terminal fitness node of seed mass was included as the cumulative seed mass for the current and previous year, for each given year of study. We followed the recommendation of Bolker *et al* (2009) and Geyer *et al* (2013) and treated block as a fixed effect. Thus, we avoid the computational issues associated with the relaxation of Gaussian assumptions for random effects that relate to all generalized linear models, including aster models, and are explicitly motivated by the need to model cases that do not meet these specific assumptions. For each region and year combination, mean fitness was represented by the median block estimate for fitness (and standard error).

**Figure 2.**
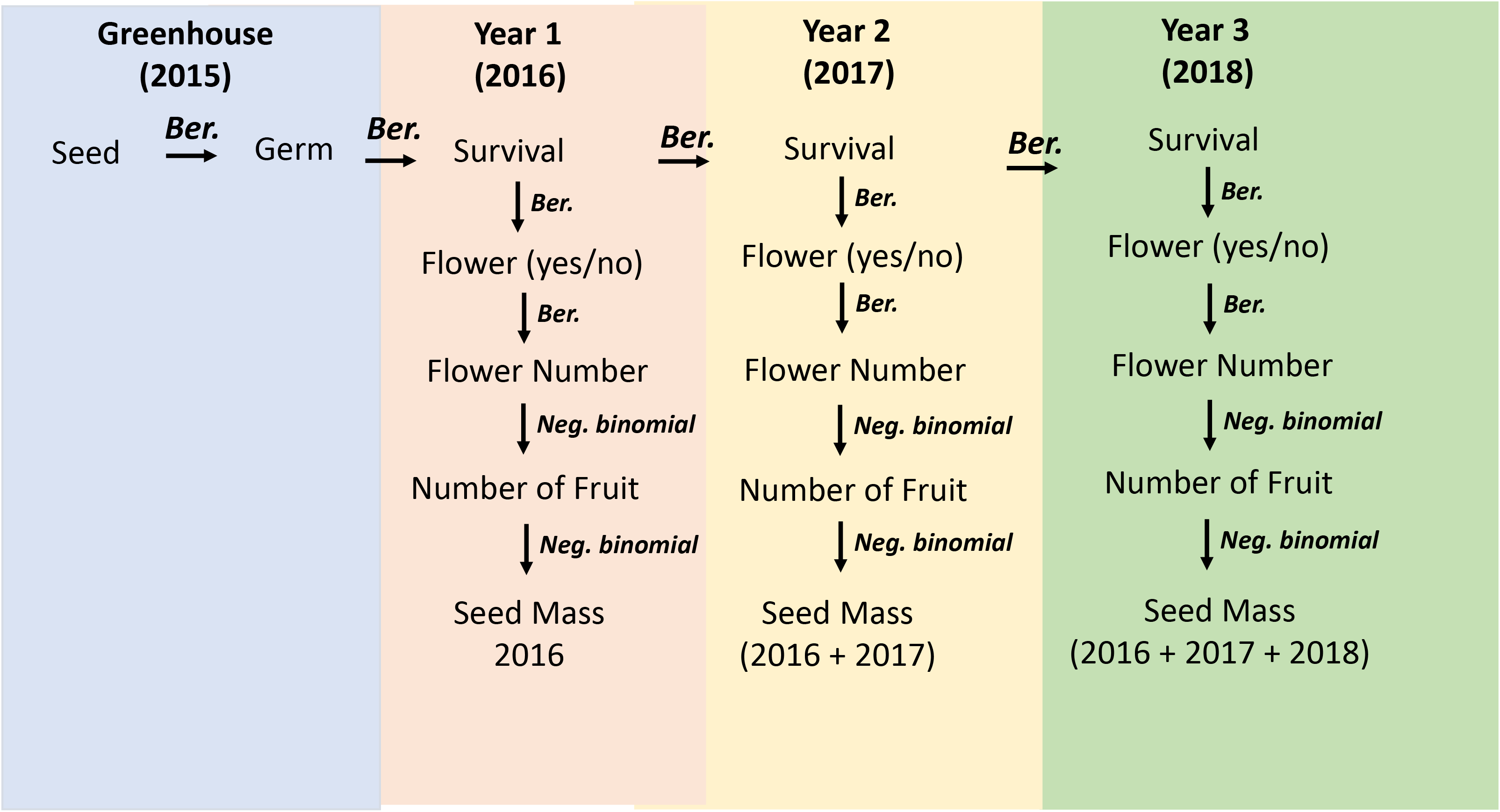
Graphical model used to estimate lifetime fitness for each plant in the common garden. Each node represents a fitness component and therefore response variable, and arrows represent conditional distributions. Probability of germination, flowering, and survival (0 or 1; Bernoulli distribution), and total number of flowers, total number of fruits and seed mass (negative binomial). Seed mass for each year was calculated as the sum of seed mass for the current year and all previous years of the experiment.

Fitness landscapes, characterized with aster, were calculated to describe selection on individuals in the common garden, while accounting for historical selection in source populations through the addition of population-specific environmental summaries. To analyze the expression of early phenological transitions (time to emergence in 2016) and life history trait variation (number of flowers produced in each year) in the common garden environment, year-specific aster models were used to calculate cumulative fitness landscapes. Time to emergence and annual variation in flower production were assessed as they exhibited a relatively high proportion of heritable variation (Table 2), with increased potential to respond to selection. Selection associated with the environment of origin for each population was included in the aster fitness landscape models using the first principal component that differentiated population climate as established in Yoko et al. (2020) based on 26 average annual climate variables estimated from ClimateNA (Wang *et al*, 2016). We estimated selection across populations planted within the common garden using the first axis of climatic variation, PC1 (43% of total variation) which previous research suggests largely follows a gradient in seasonal water availability, and a second axis of climatic variation, PC2 (~27% of total variation) that largely follows a temperature gradient (Yoko *et al*, 2020). To allow for correlational selection in fitness landscapes, aster models included the cross-products between the traits (days to emergence or number of flowers) and the first principal component following Geyer and Shaw (2010). For each fitness landscape, a summary of climatic variation (PC1) of origin was used as a predictor variable for yearly trait observations within the common garden, including days to emergence (2016) and annual number of flowers produced (2016-2018). Selection was represented by distinct fitness contours associated with the year of trait observation. The steepness of the fitness landscape topography, that is the magnitude of selection, is reflected by the proximity and the increment of change between contour lines. We superimposed observed individual-plant phenotypes on fitness landscapes to show the distribution of individuals sourced from each region within the estimated selection surfaces.

## Results

### General patterns of fitness expression and phenology

Plants sourced from the two alvar habitats exhibited similar initial success in emergence relative to plants sourced from prairie habitats. The proportion of seedlings that emerged from planted Great Lakes alvar (972 of 1312 = 0.741) and Manitoba alvar populations (180 of 239 = 0.753) were greater than seedlings planted from Prairie populations (298 of 790 = 0.377). Similar differentiation between alvar and prairie regions were observed for early phenological expression: the mean number of days between planting to emergence (Great Lakes alvars: 10.8, Manitoba alvars: 11.7, and Prairies: 14.5, all *t* > 3.38, P < 0.001), planting to the production of true leaves (Great Lakes alvars: 18.4, Manitoba alvars: 18.9, and Prairies: 21.5, all *t* > 1.99, P < 0.05), and days to first flower (Great Lakes alvars: 265.2, Manitoba alvars: 271.1, and Prairies: 266.1, all *t* < 0.85, P > 0.390). Such regional consistency implies a common heritable basis for early life history transitions in extreme, but predictable alvar habitats relative to extreme, but unpredictable prairie habitats. Interestingly, early similarities between populations sourced from different alvar regions disappeared with later expressions of phenology. For example, the mean number of flowers produced in the second year of the study (when more plants flowered to permit comparison) diverged based on habitat origin (Great Lakes alvar 8.8, Manitoba alvars 4.4, Prairie 3.1, all *t* < −8.87, P < 0.0001). Finally, total mean seed mass (accounting for differing numbers of plants per region) differed widely across all three regions (Great Lakes alvar 898.6 mg, Manitoba alvar 234.9 mg, Prairie 138.5 mg, all *t* < −6.43, P < 0.0001).

### Narrow-sense Heritability and Evolvability: Fitness Expressions and Phenology

Narrow-sense heritabilities for fitness expressions (Table 2A) were generally lower than those estimated for phenological traits (Table 3A). For fitness expressions, heritability ranged from 0 to 0.317, with heritability of fitness expressions from earlier life-history stages greater relative to later life-history fitness expressions. Similarly, the heritability for phenological traits ranged from 0 to 0.202, with a greater contribution of heritable genetic variation to early phenological transitions relative to later phenological transitions. Keeping with these estimates, evolvabilities (coefficient of additive genetic variation) were also generally greater for early life-history fitness expressions (Table 3A) but were generally non-existent for phenological traits (Table 3B). Moreover, and in accordance with patterns of early phenological traits, the estimates of narrow-sense heritability for the number of days to emergence was relatively consistent between the two alvar regions (Great Lakes: 0.257, Manitoba: 0.154) when compared to populations from the prairie region (0.026). Only two individuals from the Manitoba alvar and Prairie habitat types produced fruit in 2016. Therefore, we did not attempt to estimate heritability or evolvabilities for fruit set or seed mass in 2016 from these two regions.

**Table 3.** (A) Narrow-sense heritability and standard error and (B) evolvability (coefficient of additive genetic variation) of *Geum triflorum* phenology events grown in a common garden.

| Region | Number of Days to Emergence | Planting to First Flower 2016 | Planting to First Flower 2017 | Planting to First Flower 2018 | Planting to Fruit 2017 | Planting to Fruit 2018 | Planting to Bolt 2017 | Planting to Bolt 2018 |
| --- | --- | --- | --- | --- | --- | --- | --- | --- |
| Great Lakes | 0.257 | 0.002 | 0.202 | 0.197 | 0.000 | 0.059 | 0.059 | 0.011 |
| Alvar | (0.021) | (0.001) | (0.003) | (0.057) | (0.000) | (0.018) | (0.045) | (0.012) |
| Manitoba | 0.154 | 0.000 | 0.136 | 0.086 | 0.010 | 0.013 | 0.001 | 0.013 |
| Alvar | (0.070) | (0.000) | (0.054) | (0.130) | (0.042) | (0.026) | (0.015) | (0.037) |
| Prairie | 0.026 | 0.000 | 0.000 | 0.000 | 0.000 | 0.013 | 0.001 | 0.001 |
|  | (0.016) | (0.000) | (0.000) | (0.000) | (0.000) | (0.021) | (0.012) | (0.008) |

B
| Region | Number of Days to Emergence | Planting to First Flower 2016 | Planting to First Flower 2017 | Planting to First Flower 2018 | Planting to Fruit 2017 | Planting to Fruit 2018 | Planting to Bolt 2017 | Planting to Bolt 2018 |
| --- | --- | --- | --- | --- | --- | --- | --- | --- |
| Great Lakes | 0.111 | 0 | 0 | 0.003 | 0 | 0.001 | 0.004 | 0.001 |
| Alvar |  |  |  |  |  |  |  |  |
| Manitoba | 0.147 | 0 | 0.002 | 0.003 | 0.001 | 0.001 | 0.001 | 0.001 |
| Alvar |  |  |  |  |  |  |  |  |
| Prairie | 0.049 | 0 | 0 | 0 | 0 | 0.001 | 0 | 0.001 |

More generally, heritability estimates for fitness expressions and phenological traits exhibited consistent variation among habitat types. Populations from both alvar regions (Great Lakes and Manitoba alvars) expressed a larger proportion of heritability relative to populations from the prairie region (Table 2).

### Mean Fitness and Selection

Mean fitness, determined as annual cumulative seed mass, varied widely across regions and years. Thus, we performed an analysis of regional differences across the course of each year of study as the effect of region, year and their interaction where all were significant predictors for fitness (all test deviance > 82.09, all P < 0.0001). Fitness represented by the median block estimate (and standard error) was consistently higher in plants originating from the Great Lakes alvar region for all three years of study. This contrasted with plants from the Manitoba alvar and Prairie regions which produced almost no seed until the second year of study (Fig. 3), precluding fitness comparisons among regions in 2016. Plants from prairie populations consistently exhibited reduced fitness relative to Great Lakes alvar populations, with Manitoba alvar plants intermediate to the two regions. Region-specific estimates of fitness increased throughout the course of our study. Whereas the rate of yearly fitness increase was much greater in Great Lake compared to Manitoba alvars in the first two years of our study, the rate of annual fitness increase began to converge across regions in the third year.

**Figure 3.**
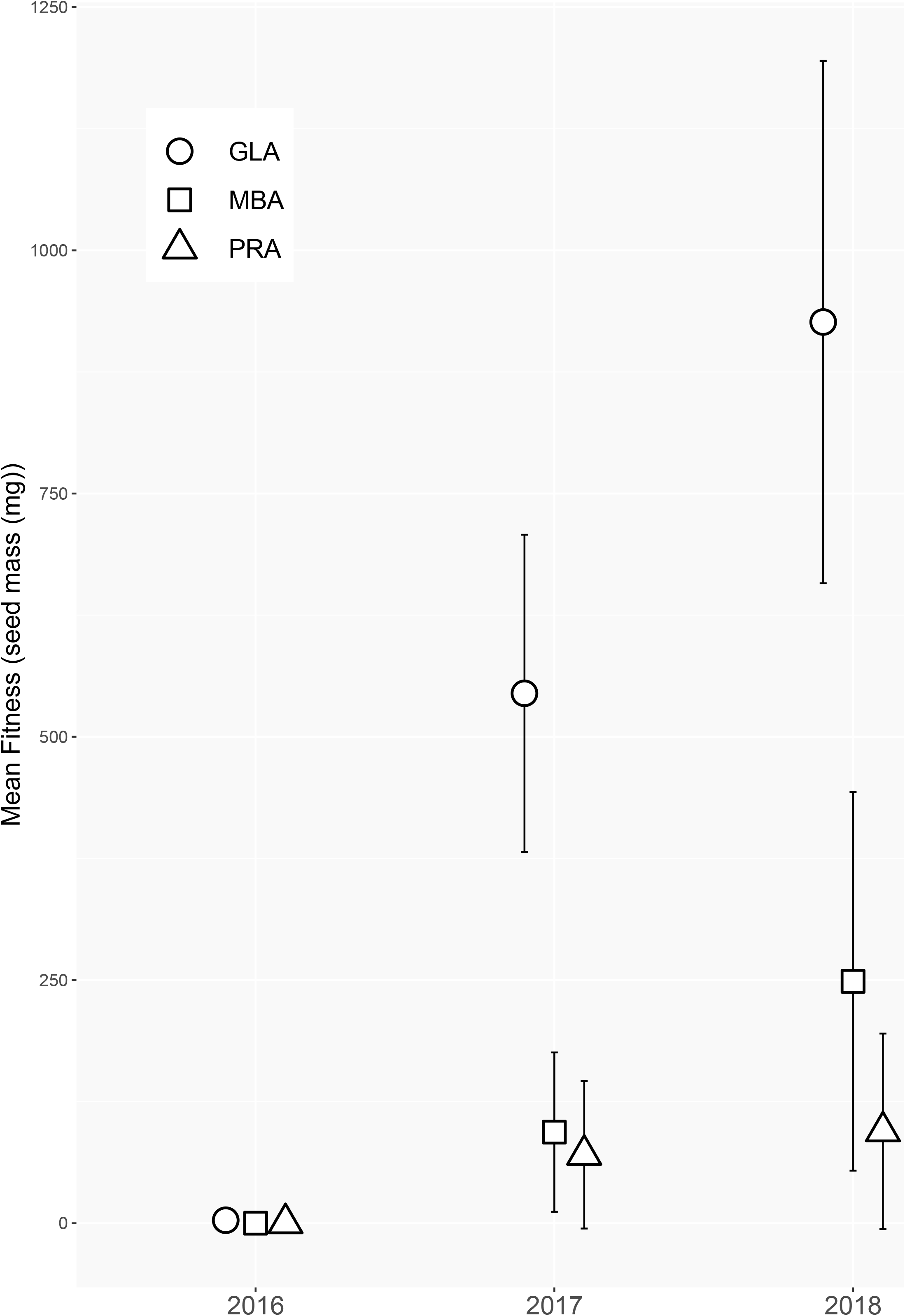
Estimated mean fitness and standard errors for plants from Great Lakes Alvars (open circles), Manitoba Alvars (open squares), and Prairie (open triangles) source populations across three successive years.

Calculation of fitness landscapes identified fitness optima for days to emergence in 2016 (supplementary Figure S1) and number of flowers for all three years of the study (Fig. 4A-4C) with a principal component of source-population environmental variation. This principal component primarily described a soil moisture gradient associated with population origin (Yoko et al. 2020). Regardless of trait-PC1 combination, selection was weak in 2016 with small fitness changes across fitness intervals. In subsequent years, selection on flower number and PC1 became stronger, and the range of optimal flower number-PC1 combination became successively narrower across years (Fig. 4). Consistent with estimates of mean fitness, individuals from the Great Lakes regions were consistently distributed closest to the fitness optima in each year of study.

**Figure 4.**
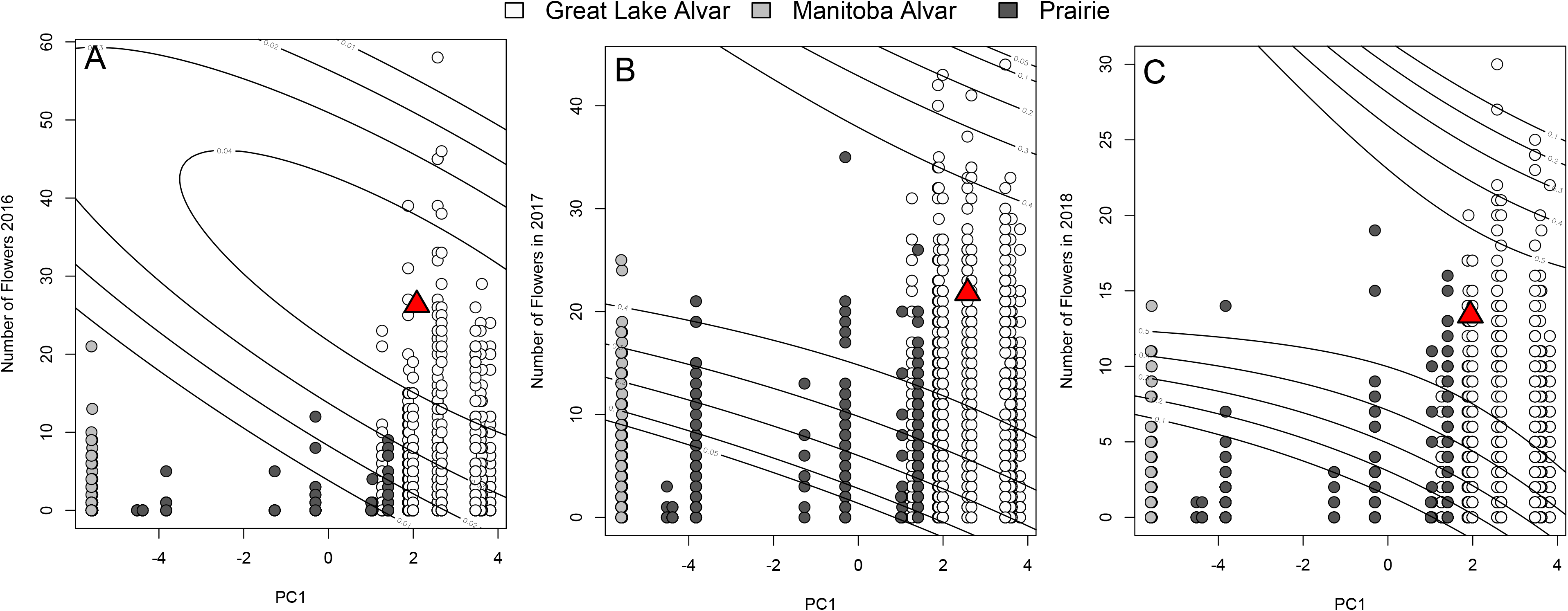
Fitness landscapes for the total number of flowers produces and a principal component describing a moisture gradient based on climate of origin for seed source populations from Yoko et al. (2020) over three consecutive years in the common garden (panel A-C). Points denote observed numbers of flowers, and line contours indicate fitness contours determined with aster models following Geyer and Shaw (2008). Great Lakes Alvars (open circles), Manitoba Alvars (light grey circles), and Prairie (dark grey circles) source populations and identified fitness optimum (red triangle) are indicated.

## Discussion

Optimizing fitness across coordinated life history events requires synchronous phenotypic responses to environmental cues. This challenge is made more complex when seasonal cues are unpredictable and are expected to become less predictable under global change. In this study, we show that predictable seasonal cues in alvar environments have led to substantial genetic control over early life history traits, such as probability and days to emergence, but plasticity in later life history traits enable longer term fitness tracking. In contrast, individuals from prairie habitats, which are characterized by less predictable seasonal cues, exhibit overall reduced heritability for the same traits. This reduced heritability may limit the ability of a perennial species to traverse a fitness landscape across generations. Thus, variance in the genetic or environmental contribution to phenological and life-history trait variation may have substantial influence on fitness, particularly when considering the capacity for individuals to maintain fitness across environments as seasonal cues shift under global change.

### Heritability and Evolvability in the Common Garden

The potential for traits to respond to selection is in part dependent on the degree of standing genetic variation for traits under selection. Individuals of *G. triflorum* from predictable alvar habitats exhibited overall larger estimates of narrow-sense heritability in both fitness expressions (Table 2) and phenology (Table 3) compared to plants from unpredictable prairie habitats. In particular, the probability of emergence and total number of flowers produced in the common garden, features important to fitness, were higher for alvar compared to prairie plants. Further, heritability of the phenological aspect of emergence (number of days to emergence, Table 3) was 1.5 to 2 orders of magnitude greater in plants from alvars compared to prairie habitats. Importantly, this has implications for estimates of evolvability as the timing of emergence for plants from alvar habitats were twice that of plants from prairie habitats. Therefore, the probability and timing of emergence is expected to be approximately twice as responsive to selection in plants from alvars compared to prairies. Interestingly, the probability of emergence, that exhibits the highest degree of evolvability, was comparable across regions (Table 2), suggesting that although the heritability for emergence in prairie environments is reduced, the potential response to selection is similar across habitat of origin.

#### Genetic and environmental contributions to phenology

The timing of seedling emergence is an ecologically important aspect of phenology and closely associated with fitness. When a seedling emerges determines the future environment experienced during growth, reproduction, and seed dispersal (Evans and Cabin, 1995), and has thus been interpreted as a form of niche construction (Donohue, 2005). Therefore, local adaptation should include a strong heritable basis for the timing of emergence. In our study, estimates of heritability and evolvability were greater in plants from alvar compared to prairie environments. With predictable seasonal extremes of flooding and drought in alvars (Reschke *et al*, 1999), individuals have little time to complete reproductive cycles. Therefore, a strong heritable basis for the timing of emergence should be prevalent in alvar environments to maximize the amount of time for reproductive effort before the onset of late-season dormancy. In contrast, prairie environments lack this consistent signal of environmental coordination and therefore employ a more attenuated emergence schedule. Greater plasticity in timing of emergence is reflected in overall lower estimates of heritability in plants from PRA populations (Table 3).

An alternative explanation for the difference in the magnitude of heritability between GLA and PRA plants could be attributed to variation in the demography of GLA populations. These populations are generally large and exhibit increased density within similar spatial extents relative to prairie populations (Hamilton and Eckert, 2007). Further, alvar habitats typically experience reduced competition due to the unique environmental features that support the persistence of select flora on these habitats (Partel *et al*, 1998). This contrasts with prairie habitats where PRA populations may also be fragmented, but forbs often experience increased competition with native and invasive grasses (Dickson and Busby, 2009).

#### Consequences of predictable and unpredictable environments

Predictably varying environments present populations with a consistent pattern of selection, resulting in an evolutionary response provided sufficient standing genetic variation exists. However, consistent and predictable selection need not necessarily erode additive genetic variation for traits as traditionally conjectured for traits presumed to be closely associated with fitness (Roff and Mousseau, 1987; McFarlane *et al*, 2014). Provided that sufficient annual variation in the predictable environmental cue exists, optimal responses to this cue will vary across years. Therefore, fluctuation in the direction and magnitude of selection may maintain appreciable additive genetic variation (Bell, 2010). Such a pattern was observed in the timing of emergence and flowering regardless of habitat of origin (Table 3).

Given the importance of the timing of emergence to subsequent life-history events, a strong genetic basis could provide a consistent start point for life cycles. However, greater environmental variation associated with later life history events could cumulatively impact the evolutionary trajectory of phenotypic traits. Indeed, later life-history expressions have been predicted to exhibit relatively reduced estimates of heritability (Price and Schluter, 1991). Therefore, the degree of plasticity for post-emergence life-history events would be much greater than during early life-cycle events. Estimates of heritability and evolvability for the timing of emergence were always greater regardless of habitat of origin. Heritability for the timing of flowering was reduced, but still appreciable compared to the remaining life history events. This pattern matches our predictions of a greater importance of heritable variation in early life history events with a minor role for plasticity. During later events, environmental variance is greatest and therefore plasticity will have a relatively larger effect on phenotypic variance. This pattern implies some continuum of the relative effects of heritable and environmental variance on phenotypic variation. Such a continuum implies an intermediate life-history expression that has an approximately equal proportion of heritable and plastic determinism. Given the intermediate estimates of heritability for the timing of flowering (across years; Table 3), this life-history event may represent such an intermediate. Indeed, this phenological signpost has been shown to exhibit similar magnitudes of narrow-sense heritability in similar systems (Burgess *et al*, 2007; Geber and Griffen, 2003; Goncalves-Vidigal *et al*, 2008; Mitchell and Shaw, 1993; O’Neil, 1997; Wheelwright, 1985), and other studies confirming adaptive plasticity in the timing of flowering (Donohue *et al*, 2000; Jimenez-Ambriz *et al*, 2007). Plasticity in flowering time seems to be more common than not (Levin, 2009). For example, Ensing and Eckert (2019) report a gradient of plasticity that resulted in the timing of flowering in *Rhinanthus minor* transplanted along a range of altitudes that plastically shifted flowering time to match those of new conspecifics. This midpoint life history event may represent the transition when environmental variation begins to exceed heritable variation in determining phenotypic variation, and ultimately population fitness.

#### Interannual patterns of heritability

Estimates of heritability can vary across years for the same trait in the same locality. However, if heritability estimates are constant across environments some ability to predict phenotypes as a response to selection exists. For example, Young *et al* (1994) determined consistent heritability of floral traits in *Raphanus sativus* in three different environments, indicating the absence of a genotype-by-environment interaction. Therefore, the expectation is that selection acting on these traits would result in environment-specific changes in phenotypes. In contrast, our results suggest that in general, heritability for expressions of fitness (flower number, fruit number, and seed mass) of plants from alvar habitats decreased over time, supporting the prediction from Price and Schluter (1991) that the cumulative exposure to environmental variability limits the estimate of narrow-sense heritability. The same traits in plants from prairie environments exhibited very little variation across years, and overall negligible estimates of heritability. Similarly, heritability estimates for fitness expressions were overall smaller than those for phenology traits, regardless of habitat origin.

The expression of additive genetic variation is dependent on local environmental conditions (Hoffmann and Merila, 1999; Sheth *et al*, 2018) and may change under unfavorable or stressful conditions (Emery and Ackerly, 2014; Schlichting, 2008). Therefore, the potential rate of the response to selection will depend on the environment-specific expression of additive genetic variation. For example, Torres-Martinez *et al* (2019) found greater short-term potential for adaptation during stressful drought (La Niña) years compared to more favorable wet (El Niño) years in an experimental precipitation gradient in *Lasthenia fremontii*. The temporal availability of water in alvar habitats likely imposes severe drought stress that accompanies periods of drought (Lundholm and Larson, 2003; Rosén, 1995; Schaefer and Larson, 1997). Individuals in our study from alvar communities likely experienced drought stress during later life history stages, predicting higher expressions of additive genetic variance and narrow-sense heritability (but see Blows and Sokolowski, 1995; Charmantier and Garant, 2005). This may seem in conflict with the above discussion of later life-history events exhibiting smaller estimates of heritability, as found in our study. However, enhanced expression of additive genetic variance does not necessarily equate to larger estimates of heritability, but rather, depends on the proportion of environmental variation associated with traits. Therefore, the cumulatively larger environmental variance associated with the later life-history events occurring throughout times of drought could reduce heritabilities regardless of enhanced expression of additive genetic variance. Our estimates of low heritability and evolvability of life-history events during these periods of stress suggest that environmental variance could hamper an adaptive response to selection (Falconer and Mackay, 1996; Levin, 1998, but see Ghalambor *et al* 2007).

### Mean Lifetime Fitness and Fitness Landscapes

#### Mean Lifetime Fitness

Mean fitness, as determined through aster models, relates directly to per capita rates of population increase, and therefore population sustainability and growth. Interestingly, plants from GLA populations consistently exhibited the highest fitness estimates in the common garden environment across all three years (Fig. 3). This was unexpected as alvar habitats differ markedly, especially in terms of water availability, from prairie habitats. Even plants sourced from prairie environments in close proximity to the common garden did not perform nearly as well as plants from either MBA or GLA populations. The large discrepancy in fitness may be attributed to low germination success in prairie plants (37.7%) compared to plants from Great Lakes (74.1%) and Manitoba (75.3%) alvars. Overall, fitness was low in the first year of our study, as plants became established in the common garden and with relatively fewer plants flowering compared to subsequent years. In the remaining two years of study, as fitness increased across all regions of origin, the difference in fitness among regions increased while maintaining the same pattern of fitness expression (Fig. 3). Therefore, the pattern of plants from GLA populations exhibiting higher fitness in the common garden was not a short-term artifact of establishment.

The expression of fitness, much like the expression of additive genetic effects, is dependent on the environment. When genotypes are moved from their home range to a novel environment, fitness may decrease as a sign of local adaptation (Garrido *et al*, 2012; Hereford, 2009), increase (e.g., Sheth et al. 2018), or remain constant (Galloway and Fenster, 2000). In our study, the movement of genotypes that evolved under GLA environments to the prairie environment of the common garden resulted in an increase in fitness. Yoko et al. (2020) detected trait enhancements associated with water-use efficiency, among others, in plants from alvars compared to prairie environments. Plants originating from alvars would experience greater water availability in the prairie environment of the common garden, where thick rich soils mitigate unpredictable fluctuations in water availability (Risser *et al*, 1981;Anderson, 2006). Therefore, enhanced water-use efficiency of alvar plants could provide a physiological advantage over prairie plants that would not historically experience predictable seasonal drought conditions. Finally, ecological differences rather than physical distances are likely more important in determining fitness in our common garden. However, to fully evaluate the preadaptation of alvar plants would require a reciprocal transplant experiment with both alvar and prairie populations.

#### Fitness Landscapes

Annual changes in mean fitness across source habitats corresponded with changes in the magnitude of selection as determined through fitness landscapes. Total flower number is commonly found to be under selection (Harder and Johnson, 2009) and therefore closely linked to fitness. We found moderate selection on the total number of flowers produced along an axis of environmental variation (moisture gradient) for source populations. The magnitude of selection along the axis of environmental variation represents the difference in environment between source populations and the common garden, describing the discrepancy between fitness optima across source populations and the novel common garden environment. Thus, the change in selection along the environmental gradients represents the degree of maladaptation following introduction to a novel environment.

The location of fitness optima consistently tracked mean fitness for each source population habitat type (Fig. 3, Fig. 4). Fitness across all source populations were modest in the first year of study with few individuals reproducing. Consequently, the fitness landscape (Fig. 4A) was relatively flat with a wide but shallow plateau surrounding the optima. In the two remaining years of the study, the topography of the landscapes became steeper indicating stronger selection, with successively more plants occupying the region around the fitness optima. Regardless of year, plants from the Great Lakes alvars were consistently closer to the fitness optima. Alvar habitats experience consistent extremes in water availability with annual late-season desiccation (Catling and Brownell, 1995). Lack of water availability favors selection for plants with reduced water potential and enhanced water-use efficiency (Craine and Dybzinski, 2013). Thus, plants originating from predictably water-stressed environments like alvars, could express release from water-use constraints, effectively enhancing fitness in response to increased water availability within the prairie common garden environment. In contrast, plants from prairie populations did not experience such a drastic change in water availability. Therefore, and unexpectedly, the degree of environmental maladaptation in the common garden environment was greater for plants from prairie populations than for plants from alvar populations.

Over successive years, the breadth of the fitness optimum increased along the axis of environmental variation but was relatively constrained along the axis of total flower production. Given the relatively high heritability in flower number across all three years of study (Table 2), the limited plasticity in flower number may be responsible for this constraint. In contrast, enhanced plasticity for water use traits has been associated with improved survival and seed production (Nicotra and Davidson, 2010). Therefore, the more predictable seasonal changes in water availability in alvar habitats may promote enhanced plasticity in water use traits, whereas flower number would remain relatively constant within a habitat type. However, across habitat types, total flower production was greater in plants from alvars with predictable changes in water availability compared to prairie habitats. Regardless of habitat origin, the degree of plasticity associated with flower number was restricted compared with that of water availability. Therefore, the restricted plastic response of flower production would impose a strong limitation to fitness and local adaption in the common garden environment.

Total fitness is the cumulative expression of an individual’s fitness components across its life cycle, from emergence and survival to the total number of offspring produced. Individual fitness expressions are the results of genetic and environmental effects, as well as their interaction. We observed that the independent and interactive effects of these components vary continuously across life histories. The relative degree of genetic and environmental contribution to trait variation provides a new perspective to the dynamics of life-history traits. We found a shift from genetic to greater environmental effects across successive years, allowing plants furthest from the fitness optimum to traverse the fitness landscape and increase proximity to the fitness optimum. Plants sourced from predictable alvar environments started closest to, and maintained proximity, to the fitness optimum across years. Whereas plants from unpredictable prairie environments were initially furthest from the fitness optimum, these plants were able to quickly approach the fitness optimum. Overall, these results highlight the importance of environmental variation in facilitating movement across the fitness landscape, and genetic variation in maximizing fitness near the optimum.

## Acknowledgements

The authors thank Jon Sweetman, Chad Stratilo, Mary Vetter, Rebekah Neufeld, Tyler Stadel, Steve Travers, and the Nature Conservancy of Canada for help with initial field sampling for seeds. In addition, we thank Stephen Johnson, Nick Hugo, Alexis Pearson, Zoe Portlas, Naomi Hegwood, Storm Nies, and Kate Volk for assistance in the field. Thanks also to Tony Bormann (MSUM Science Center) for logistical support and Ruth Shaw, Frank Shaw, and Charles Geyer for guidance with aster modeling.

This work was supported by a new faculty award from the office of the North Dakota Experimental Program to Stimulate Competitive Research (ND-EPSCoR NSF-IIA-1355466) and funding from the NDSU Environmental and Conservation Sciences Graduate Program to J.A.H.

## Author Contributions

JH led the initial field collections and designed the study. JH and ZY participated in field work and data collection, MWK performed the main analyses. MWK led the manuscript writing with considerable input from JH and ZY.

## Data Accessibility

All data and R code have been archived at: https://tinyurl.com/82rppcte

Supplemental Figure A. Fitness landscapes for the number of days to emergence from planting and a principal component describing a moisture gradient in source populations from Yoko et al. (2020) for the first year of study (2016). Points denote observed numbers of days to emergence and line contours indicate fitness contours determined with aster models following Geyer and Shaw (2008). Great Lakes Alvars (open circles), Manitoba Alvars (light grey circles), and Prairie (dark grey circles) source populations and identified fitness optimum (red triangle) are indicated.

