## Supplementary figures and images for "Chasing the fitness optimum: temporal variation in the genetic and environmental expression of life-history traits for a perennial plant"

### Supplemental Figure 1

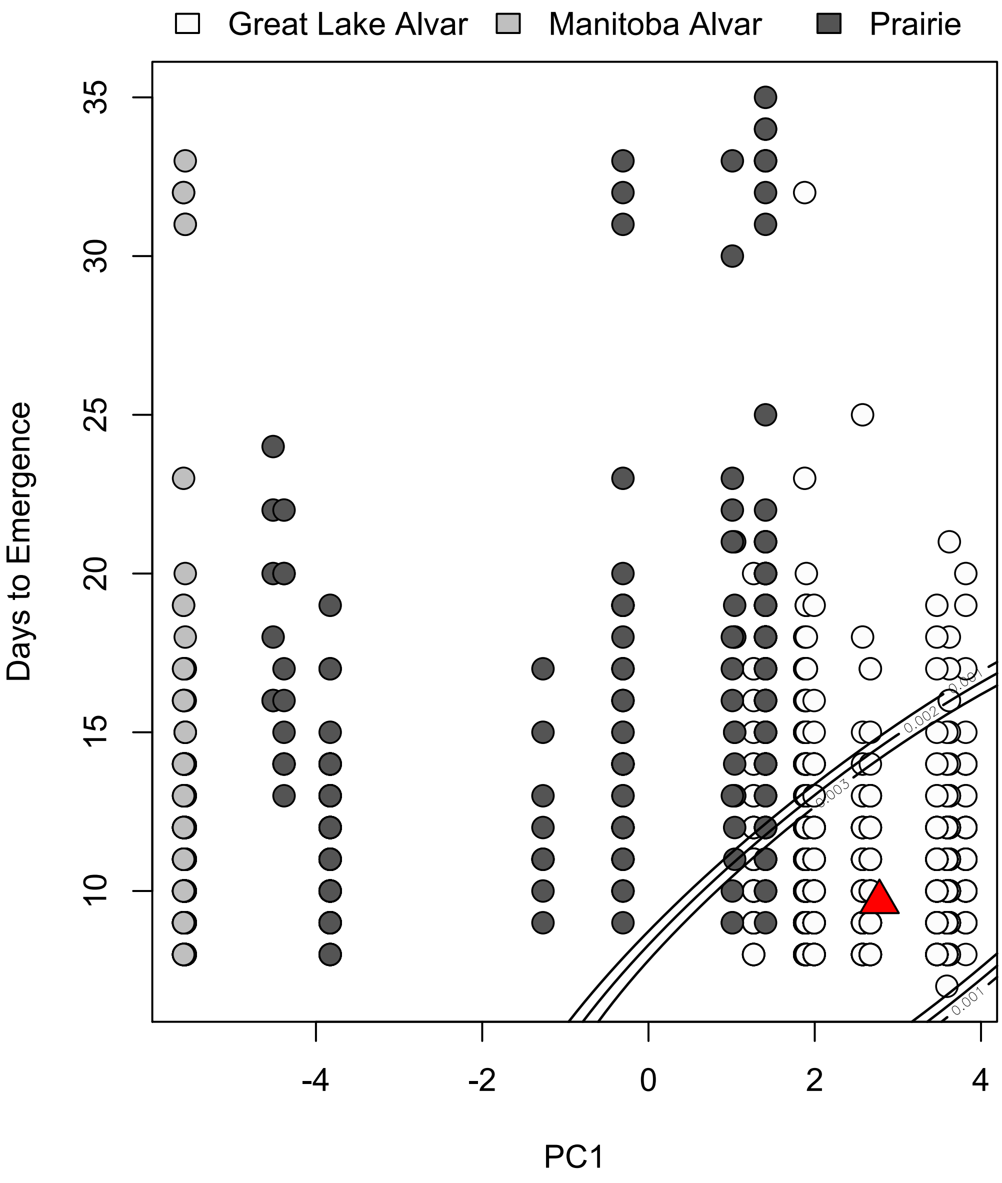
